# Novel African Rhinolophus bat ACE2 sequences reveal the determinants of Afro-Eurasian sarbecovirus entry

**DOI:** 10.64898/2026.04.02.716099

**Authors:** Yueying Zhang, Shigeru Fujita, Masahiro Kajihara, Katendi Changula, Bernard Mudenda Hang’ombe, Shusuke Kawakubo, Spyros Lytras, Jumpei Ito, Akinori Kanai, Yutaka Suzuki, Ayato Takada, Kei Sato

## Abstract

Sarbecoviruses, including SARS-CoV and SARS-CoV-2, are frequently linked to *Rhinolophus* bats as their putative natural reservoirs. Angiotensin-converting enzyme 2 (ACE2), a host carboxypeptidase widely expressed in mammalian tissues, plays a critical role in sarbecovirus infection by serving as the cellular receptor for the viral spike (S) protein. Given recent human outbreaks and pandemics caused by members of sarbecoviruses, and the wide distribution of *Rhinolophus* bats, it is essential to maintain surveillance of these viruses while improving our understanding of their interactions with bat hosts, particularly the ACE2 receptor. However, while *Rhinolophus* bats from Asia have been relatively well studied, African *Rhinolophus* bats remain underrepresented and require further investigation. In this study, five *Rhinolophus* bat lung samples were obtained from Zambia, and ACE2 genes from these individuals were cloned and sequenced. We further evaluated the susceptibility of ACE2 variants to a panel of sarbecoviruses, revealing key residues that influence viral infectivity. ACE2 polymorphism was observed among *Rhinolophus simulator* individuals, revealing multiple ACE2 genotypes within the sampled population. However, *R. simulator* ACE2s did not permit infection by the clade 3 Afro-Eurasian sarbecoviruses tested in this study. Notably, RhGB01 and BM48-31 virus utilized only *Rhinolophus blasii* ACE2. Mutational analyses further suggested that ACE2 residues 31 and 41 play important roles in modulating spike-ACE2 interactions. This study reports 4 unique ACE2 sequences of *R. simulator* and *R. blasii*, and provides new insights into the molecular interactions between African *Rhinolophus* species ACE2s and the S protein of sarbecoviruses circulating in Africa and Europe.

**Importance:** As putative natural reservoirs of sarbecoviruses, including SARS-CoV and SARS-CoV-2, *Rhinolophus* bats play a critical role in the emergence of zoonotic coronaviruses, making it essential to understand their interactions with these viruses for future pandemic preparedness. While Asian *Rhinolophus* bats have been relatively well studied, African species remain underrepresented, highlighting the need for further investigation. In this study, we cloned and sequenced ACE2 genes of five *Rhinolophus* bats collected in Zambia, Africa. We identified ACE2 polymorphism among *Rhinolophus simulator* individuals, although this variation was not associated with susceptibility to the clade 3 Afro-Eurasian sarbecoviruses examined. In addition, we identified key ACE2 residues that govern SARS-CoV-2 spike-ACE2 interactions and contribute to distinct infectivity patterns across species. These findings expand our understanding of the molecular determinants of sarbecovirus host range and support improved surveillance and risk assessment of emerging coronaviruses.

## Introduction

The COVID-19 pandemic, caused by SARS-CoV-2, has been devastating worldwide, with profound impacts on global health, economies and societies. Similarly, another virus in the same group, SARS-CoV, was responsible for the SARS outbreak back in 2003 (1). Notably, both SARS-CoV and SARS-CoV-2 belong to the *Sarbecovirus* subgenus and have been linked to *Rhinolophus* bats as their putative natural reservoirs (2–7). Members of the *Sarbecovirus* subgenus, a group within the *Betacoronavirus* genus, have been repeatedly detected in *Rhinolophus* bats (2). For instance, RaTG13 – one of the close known relative of SARS-CoV-2 - was identified in *Rhinolophus affinis* in 2013 (5, 8). These findings underscore the pandemic potential of sarbecoviruses in *Rhinolophus* bats and emphasize the urgent need to better understand the ecological and molecular interactions between these bats and the viruses they harbor.

The population of *Rhinolophus* bats is primarily distributed across the Old World, inhabit regions including Africa, Europe, Asia and Australia (9). Within the Rhinolophidae family, two major geographic clades have been identified: the Afro-eurasian and Asian clades (10). While the Oriental clade has been relatively well studied, the African clade remains insufficiently explored and lacks comprehensive investigation.

Sarbecoviruses are enveloped, positive-sense single-stranded RNA viruses. They possess a characteristic spike (S) protein that mediates host cell entry through binding to the host angiotensin-converting enzyme 2 (ACE2) receptor (11, 12). ACE2 is a physiological enzyme involved in the regulation of blood pressure (13), and is abundantly expressed in the kidney, testes, heart, and intestinal tract, but shows low or even undetectable expression in lung tissue (14). In the context of sarbecovirus infection, the receptor-binding domain (RBD) of the viral S protein binds to host ACE2 to facilitate viral entry into the cell (15). Notable members of the *Sarbecovirus* subgenus include SARS-CoV and SARS-CoV-2, both of which led to impactful outbreaks. Sarbecoviruses are categorized into clades based on their RBD sequences: clade 1a, or the SARS-CoV related clade; clade 1b, or the SARS-CoV-2 related clade; clade 2, with viruses phylogenetically close to clade 1, characterized by two deletion within receptor-binding motif (RBM); and clade 3, with viruses found in Africa and Europe, considered close to *Sarbecovirus* ancestors (15, 16). Because of the large deletions in RBM, the clade 2 viruses are unable to use ACE2 as the infection receptor (17). Clades 1a and 1b have been identified in East and Southeast Asian countries, and the receptor usage of these clades have been extensively investigated (18, 19). On the other hand, several aspects of clade 3 viruses remain under-characterized. For instance, their ACE2 usage profiles, infectivity capacity, and zoonotic potential are still largely unknown - making them important targets for further research aimed at assessing their potential for cross-species transmission.

Accumulating evidence indicates that polymorphisms in the ACE2 receptor can profoundly alter sarbecovirus infectivity by modulating S-receptor interactions. Previous studies have demonstrated that ACE2 polymorphisms in Asian *Rhinolophus affinis* bats can result in distinct viral infectivity profiles (20, 21). The binding between ACE2 and the viral S protein is highly specific, and even a single amino acid substitution can abolish infectivity. Therefore, it is crucial to further investigate the interactions between bat ACE2 and sarbecoviruses.

In this study, we cloned and sequenced the ACE2 genes from five individual African bats captured in Zambia, reporting for the first time the ACE2 sequences of the *Rhinolophus simulator* and *Rhinolophus blasii* species. We then tested the susceptibility of these newly identified ACE2 variants to a set of selected sarbecovirus S proteins using a pseudovirus infection system. Our results show that amino acid differences at key residues in ACE2 can lead to substantial variation in viral infectivity, providing new insights into the molecular basis of ACE2-S compatibility.

## Results

### *Rhinolophus* bat sbats sample collection in Zambia and species identification

To investigate the genetic diversity of African *Rhinolophus* bats sampled in this study, we first mapped the geographic locations of the bat collection sites in Zambia, a country in sub-Saharan Africa (**Figure 1A**). Both Shimabala and Suesueman Villages are located in the Lusaka Province, in close proximity to the capital city, Lusaka. Among the 5 *Rhinolophus* bat samples collected in Zambia analyzed in our study, four were obtained from Shimabala and one from Suesueman. Lung tissues were collected from all individuals for downstream molecular analyses.

**Figure 1:**
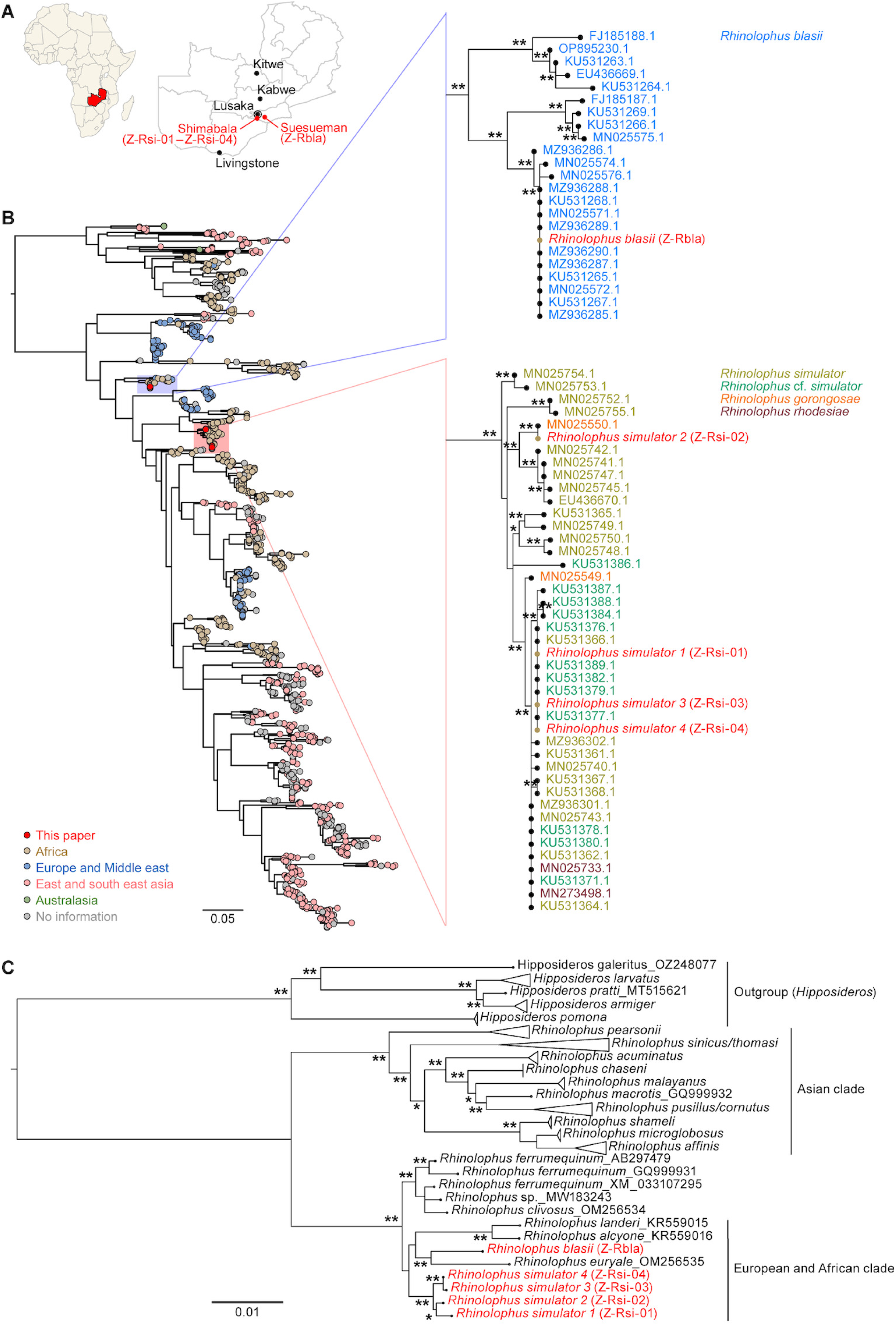
Geographic distribution and evolutionary relationship between sampled African *Rhinolophus* bats. **(A)** Geographic distribution of African *Rhinolophus* bat sampling sites in Zambia. A map of Zambia indicating major cities is shown. Field-captured *Rhinolophus* individuals are plotted according to their GPS coordinates (red dots). African map and GIS border data were downloaded from SimpleMaps.com. **(B)** Maximum likelihood phylogenetic tree of five African *Rhinolophus* bat cytochrome b (CYTB) coding sequences together with selected publicly available *Rhinolophus* CYTB sequences (Table S1). Clades containing sequences from this study are enlarged, with newly identified sequences highlighted in red. ∗, ultrafast bootstrap value >80 and ∗∗, ultrafast bootstrap value >90. Scale bar indicates genetic distance (nucleotide substitutions per site). **(C)** Maximum likelihood phylogenetic tree of five African *Rhinolophus* bat ACE2 coding sequences together with selected publicly available *Rhinolophus* ACE2 sequences (Table S1). Newly identified sequences highlighted in red. ∗, ultrafast bootstrap value >80 and ∗∗, ultrafast bootstrap value >90. Scale bar indicates genetic distance (Nucleotide substitutions per site). Country groups are highlighted in the tree based on the country data registered in the NCBI database.

We then set out to identify the species of the bat samples from Zambia. Following RNA sequencing of the lung samples, we successfully retrieved the mitochondrial cytochrome b (CYTB) sequences from all five bat individuals. A phylogenetic tree was constructed using these CYTB sequences, along with representative sequences of related *Rhinolophus* species obtained from GenBank (**Figure 1B**). The resulting tree revealed two distinct phylogenetic clusters: 4 bat samples from Shimabala were classified in the Shimabala group clustered with *Rhinolophus simulator* reference sequences, while the bat sample from Suesueman was part of the cluster of *R. blasii* sequences (**Figure 1B, right**). This analysis suggests that the 4 bats from Shimabala (hereafter referred to as Z-Rsi-01 to Z-Rsi-04) are members of the *R. simulator* species, while the one from Suesueman is a *R. blasii* (hereafter referred to as Z-Rbla) (**Figures 1A and 1B**).

To investigate the ACE2 sequences of these samples, we cloned the five bat individuals’ ACE2 from cDNA library of bat lung samples. Then, we constructed a phylogenetic tree based on their ACE2 coding sequences (**Figure 1C**). ACE2 sequences of related *Rhinolophus* species obtained from GenBank were also included for comparison. As expected, these 5 ACE2 sequences were included in the European and African clade (**Figure 1C**). Three distinct ACE2 sequences were identified among the Z-Rsi samples, indicating polymorphism of ACE2 within the population. Moreover, consistent with the CYTB-based phylogeny, Z-Rsi-01 to Z-Rsi-04 (Shimabala) formed a single clade, while Z-Rbla (Suesueman) grouped separately with *Rhinolophus euryale*, a *Rhinolophus* bat species residing in Africa. In summary, we identified five novel ACE2 sequences of the two previously uncharacterized African *Rhinolophus* species and revealed the ACE2 polymorphism among the Z-Rsi individuals.

### Different sensitivity of Zambian *R. simulator* and *R. blasii* ACE2s to sarbecovirus infection

To investigate the newly identified ACE2 usage of sarbecoviruses, we established a panel of human osteosarcoma (HOS) cells stably expressing the human TMPRSS2 protease (HOS-TMPRSS2), along with ACE2 proteins derived from the Zambian bat samples. Specifically, the ACE2 orthologs from 4 *R. simulator* (Z-Rsi-01 to -04) and 1 *Rhinolophus blasii* (Z-Rbla) were transduced into HOS-TMPRSS2 cells using lentivirus vectors to generate stably expressing lines. The expression of these 5 Zambian *Rhinolophus* bat ACE2 proteins in these cell lines was confirmed by western blotting, which showed consistent and comparable expression levels across all constructs (**Figure 2B**).

**Figure 2.**
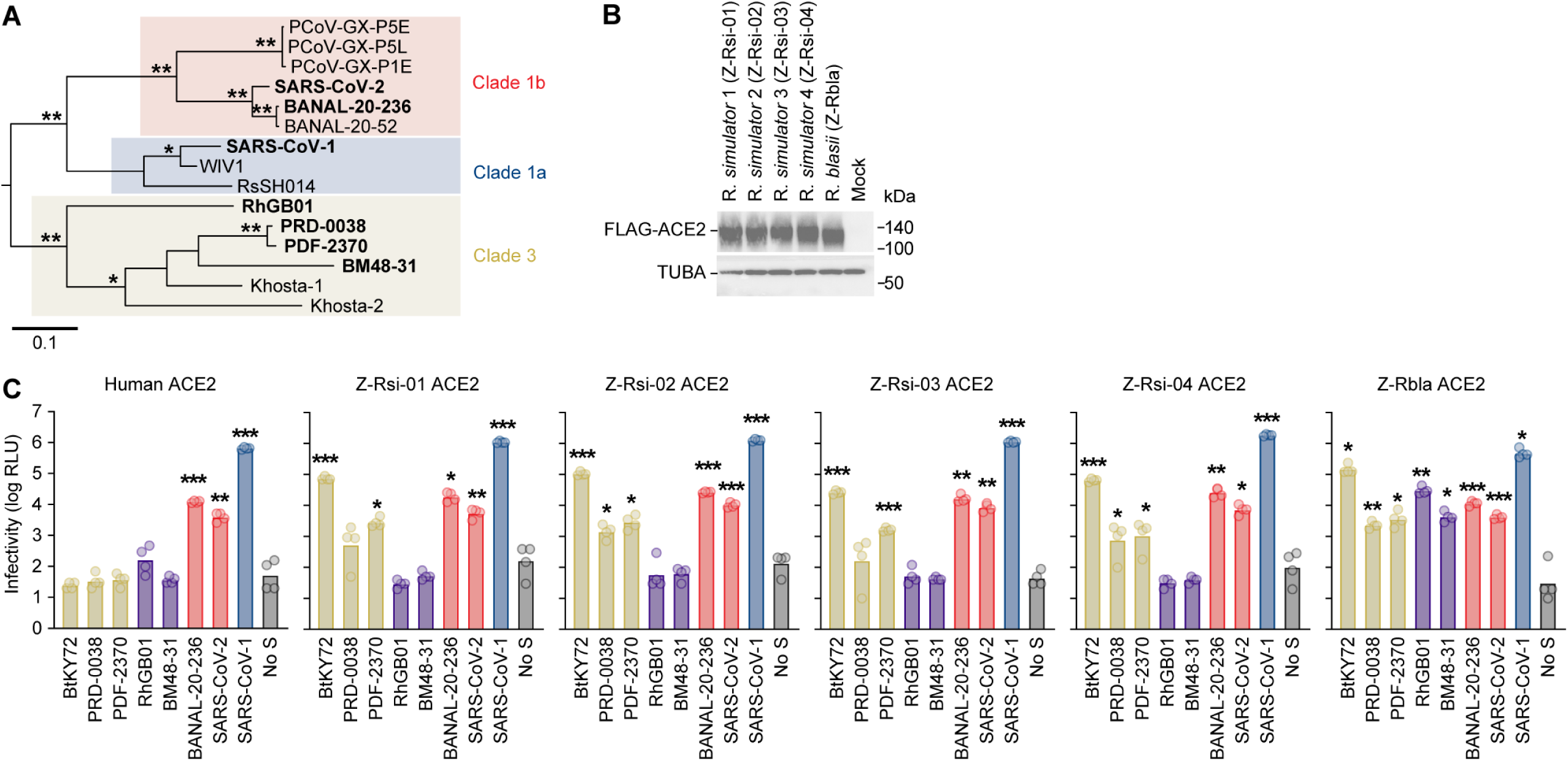
Analysis of susceptibility of 5 African ACE2 to selected sarbecoviruses. **(A)** Maximum likelihood tree of selected sarbecoviruses based on their S RBD amino acid sequences. Viruses used for pseudovirus assays are highlighted in bold. ∗, ultrafast bootstrap value >80 and ∗∗, ultrafast bootstrap value >90. Scale bar indicates genetic distance (nucleotide substitutions per site). Sarbecovirus clades are highlighted in the tree based on a previous study (42). **(B)** Western blotting. A representative blot of the HOS-TMPRSS cells stably expressing flag-tagged *Rhinolophus* bat ACE2 is shown. TUBA (α-Tubulin) is an internal control for the cells. kDA, kilodalton. **(C)** Pseudovirus assay. HIV-1-based reporter viruses pseudotyped with Spike proteins of eight sarbecoviruses were generated and inoculated into HOS-TMPRSS2 cells stably expressing *Rhinolophus* bat ACE2 at 4 ng of HIV-1 p24 antigen per well. Infectivity was quantified as relative light units (RLU) in each target cell line. Data are presented as mean ± SD from four technical replicates.

To assess viral entry efficiency, we prepared lentivirus-based pseudoviruses bearing the S proteins from eight representative sarbecoviruses. Two out of the eight are the human sarbecoviruses: SARS-CoV (strain Frankfurt 1) from clade 1a; and SARS-CoV-2 (strain Wuhan-Hu-1) from clade 1b. The other 6 are from bats: BANAL-20-236 from clade 1b; and BtKY72, PRD-0038, PDF-2370, RhGB01, and BM48-31 from clade 3, which mainly consist of European and African sarbecoviruses and corresponds with the origin of our bat samples (**Figure 2A**). We then inoculated these pseudoviruses into the HOS-TMPRSS2 cells stably expressing respective ACE2 proteins (hereafter referred to as target cells). As shown in **Figure 2C**, the cells expressing all *R. simulator* ACE2 variants (i.e., Z-Rsi-01 to -04) exhibited similar levels of susceptibility to all tested pseudoviruses: similar to human ACE2-expressing cells, all *R. simulator* ACE2 variants supported infection by pseudoviruses bearing the S proteins of clades 1a and 1b (**Figure 2C**). Consistent with previous reports (22), human ACE2-expressing cells were not susceptible to any clade 3 viruses tested (**Figure 2C**). In the case of the cells expressing all *R. simulator* ACE2 variants, they showed infectivity with three clade 3 pseudoviruses: BtKY72, PRD-0038, and PDF-2370 (**Figure 2C**). On the other hand, the other two clade 3 viruses, RhGB01 and BM48-31, did not infect the cells expressing any of *R. simulator* ACE2 variants (**Figure 2C**). These results suggest that the observed polymorphic variations within *R. simulator* ACE2 do not significantly influence the entry efficiency of sarbecoviruses tested.

In the case of the cells expressing *R. blasii* ACE2, the infection patterns of clades 1a and 1b and three clade 3 (BtKY72, PRD-0038 and PDF-2370) pseudoviruses were similar to those of *R. simulator* ACE2 (**Figure 2C**). However, it was of interest that the other 2 clade 3 pseudoviruses, BM48-31 and RhGB01, exhibited infectivity only in the cells expressing *R. blasii* ACE2 (**Figure 2C**). These results suggest that interspecies differences in ACE2, rather than intraspecies polymorphisms, may play a more critical role in determining the susceptibility of host cells to the entry of sarbecoviruses, especially at least two clade 3 viruses, BM48-31 and RhGB01.

### Residues 31 and 41 determine the infection receptor potential of *R. blasii* ACE2 for BM48-31 and RhGB01

To determine the amino acid residue(s) that are responsible for the susceptibility of *R. blasii* ACE2 to the two clade 3 viruses, BM48-31 and RhGB01, we compared the amino acid sequences of the African *Rhinolophus* ACE2 orthologs we determined in this study. Previous structural studies showed that the two amino acid residues positions at 31 and 41 of the ACE2 protein of *Rhinolophus landeri*, an African *Rhinolophus* bat, interacts with the S protein of BtKY72 (a clade 3 virus) (23). Guided by these findings, we compared the ACE2 sequences of 4 *R. simulator* and 1 *R. blasii* and identified clear differences at these two amino acid residues (**Table 1**). This observation led us to hypothesize that variations at residues 31 and 41 underlie the species-specific differences in the susceptibility to BM48-31 and RhGB01 (**Figure 2C**).

**Table 1.** Amino acid differences among African *Rhinolophus* ACE2s.

| Position | Z-Rsi-01 | Z-Rsi-02 | Z-Rsi-03/04 | Z-Rbla |
| --- | --- | --- | --- | --- |
| 21 | T | T | T | **P** |
| 31 | D | D | D | **N** |
| 34 | S | S | S | **T** |
| 35 | E | E | **Q** | E |
| 41 | Y | Y | Y | **H** |
| 173 | G | **B** | G | G |
| 216 | E | E | E | **Q** |
| 301 | N | N | N | **T** |
| 329 | N | N | N | **E** |
| 335 | D | D | D | **E** |
| 475 | K | K | K | **Q** |
| 531 | **Q** | R | R | R |
| 546 | N | N | N | **H** |
| 548 | T | T | T | **R** |
| 593 | **I** | T | T | T |
| 684 | Y | Y | Y | **F** |
| 735 | H | H | H | **Y** |
| 776 | S | S | S | **G** |

To directly assess the effect of these two residues on the pseudovirus infection of BM48-31 and RhGB01, we constructed the Flag-tagged plasmids expressing either parental *R. blasii* ACE2 or its derivatives carrying N31D, H41Y, or both substitutions (N31D/H41Y) and transfected these plasmids into HEK293 cells. Western blotting showed that the expression levels of transiently expressed ACE2 proteins were comparable (**Figure 3A**).

**Figure 3.**
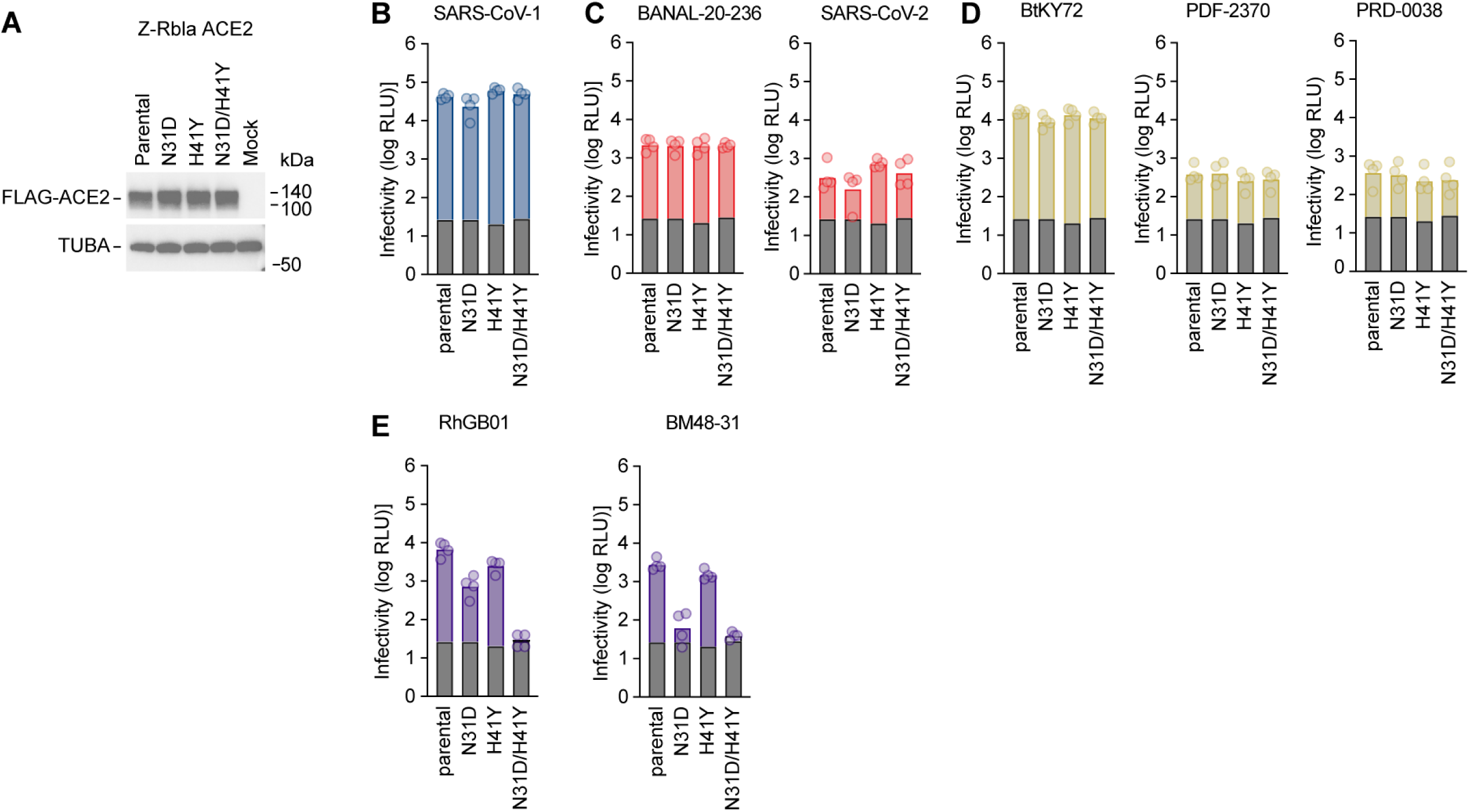
Analysis of susceptibility of *Rhinolophus blasii* ACE2 to selected sarbecoviruses. (**A)** Western blotting. A representative blot of the HEK-293T cells transiently expressing flag-tagged *Rhinolophus* bat ACE2 cells is shown. TUBA is an internal control for the cells. kDA, kilodalton. **(B-F)** Pseudovirus assay. HIV-1-based reporter viruses pseudotyped with Spike proteins of eight sarbecoviruses were generated and inoculated into HEK-293T cells transiently expressing *Rhinolophus* bat ACE2 at 4 ng of HIV-1 p24 antigen per well. Infectivity was quantified as relative light units (RLU) in each target cell line. The grey bar represents the no spike control. Data are presented as mean ± SD from four technical replicates.

We then inoculated 8 sarbecovirus pseudoviruses into these cells. As shown in **Figures 3B-3D**, the infectivity of the sarbecovirus pseudoviruses from clade 1a (SARS-CoV; **Figure 3B**), clade 2 (SARS-CoV-2 and BANAL-20-236; **Figure 3C**), and 3 clade 3 (BtKY72, PRD-0038, and PDF-2370; **Figure 3D**) was not affected by the substitutions of the amino acid residues positioned at 31 and 41. On the other hand, while the BM48-31 infectivity was comparable between parental *R. blasii* ACE2 and the N41Y ACE2 variant, BM48-31 exhibited reduced infectivity in cells expressing the N31D ACE2 variant and the ACE2 protein carrying both substitutions **(Figure 3E, left)**.These results suggest that the asparagine at position 31 plays a key role in facilitating the BM48-31 S binding to *R. blasii* ACE2. RhGB01 showed a distinct pattern from BM48-31: neither N31D nor H41Y alone substantially altered susceptibility, whereas the double substitution led to a marked loss of infectivity (**Figure 3E, right**). These observations suggest that the RhGB01 S protein likely engages both residues 31 and 41 cooperatively, and that alteration of either one alone is insufficient to disrupt the S-ACE2 interaction. Together, these results highlight residue-specific and virus-specific differences in how clade 3 sarbecovirus S proteins recognize ACE2 receptors of African *Rhinolophus* bats.

## Discussion

*Rhinolophus* bats have been recognized as natural reservoirs for several emerging viruses, including members of coronaviruses. The emergence of SARS-CoV and SARS-CoV-2—both believed to originate from *Rhinolophus* bats—highlights the critical importance of studying this bat genus (2, 7). However, limited sampling across the Afrotropical region has resulted in a constrained understanding of African *Rhinolophus* species and their ACE2 receptor genes, particularly when compared to their Asian counterparts (24). In this study, we identified and characterized novel ACE2 sequences from two African *Rhinolophus* species—*R. simulator* and *R. blasii*—captured in Zambia. Using a pseudovirus entry assay, we evaluated the susceptibility of these ACE2 variants to a panel of sarbecovirus S proteins. The results revealed differential infectivity patterns across ACE2 variants, driven by specific amino acid residues that appear to govern S-ACE2 compatibility.

Four bat individuals were captured in Shimabala and one individual in Suesueman, as shown on the map (**Figure 1A**). Both sampling sites are located in Lusaka Province, Zambia, and are relatively close to one another geographically. In the mitochondrial CYTB phylogenetic tree (**Figure 1B**), the Shimabala individuals formed a clade closely related to *R. simulator* reference sequences, supporting the classification of the Z-Rsi individuals as *R. simulator*. In contrast, the individual from Suesueman clustered with *R. blasii* sequences, indicating that this individual is most likely *R. blasii*. In the ACE2-based phylogenetic tree (**Figure 1C**), however, a slightly different pattern was observed. The Shimabala group formed a distinct clade composed exclusively of the Z-Rsi individuals, further supporting their classification as *R. simulator*. In contrast, the Suesueman individual clustered relatively more closely with the *Rhinolophus euryale* reference sequence. As shown in previous study, ACE2 evolution may not mirror neutral mitochondrial evolution (25).This discrepancy is likely due to positive selective pressures, such as host-virus coevolution, acting on ACE2.

In this study, we report for the first time the presence of ACE2 polymorphisms in African *Rhinolophus* bats. Within the *R. simulator* individuals, three distinct ACE2 genotypes were identified (**Table 1**). Similar ACE2 polymorphisms have been previously reported in *Rhinolophus affinis* populations in China and Vietnam (20, 21). We have particularly found that the ACE2 polymorphism in *Rhinolophus affinis* closely associate with the sensitivity to the sarbecoviruses identified from *Rhinolophus affinis* (20). However, unlike the case of this Asian *Rhinolophus* bat (*Rhinolophus affinis*) ACE2 and Asian sarbecoviruses, we did not observe any differences in pseudovirus infectivity among the different *R. simulator* genotypes (**Figure 2C**). This suggests that the observed polymorphisms may not affect receptor functionality, but rather reflect historical evolutionary pressure from ACE2-using pathogenic viruses. Such selective pressure may have favored the retention of certain ACE2 variants in the population. In contrast, differences in virus susceptibility were observed between the cells expressing *R. simulator* and *R. blasii* ACE2. Notably, the clade 3 sarbecoviruses RhGB01 and BM48-31, which have shown limited infectivity in other bat ACE2 systems (22), exhibited relatively high infectivity in *R. blasii* ACE2-expressing cells compared to *R. simulator* ACE2 in our assays (**Figure 2C**). This observation aligns with previous findings in our earlier study, in which *Rhinolophus alcyone*, another African *Rhinolophus* species, was the only ACE2 among 53 tested that permitted infection by both RhGB01 and BM48-31 (22). These results suggest a potential relationship between the geographical distribution of sarbecoviruses and the ACE2 compatibility of sympatric bat hosts. Furthermore, BM48-31 was originally detected in *R. blasii* in Bulgaria in 2008 (12). The susceptibility of Zambian *R. blasii* ACE2 to BM48-31 implies a conserved pattern of ACE2 usage by this virus across geographically distant populations of the same bat species.

Positions 31 and 41 in ACE2 have been identified as key determinants of sarbecovirus infectivity in multiple studies. In human ACE2, residue 31 plays a critical role in forming a hydrogen bond with the SARS-CoV-2 S protein (26). In our previous work, residue 31 was also shown to be closely associated with the host tropism of BANAL-20-52 and BANAL-20-236 (25). Additionally, another study demonstrated that residues 31 and 41 in *Rhinolophus ferrumequinum* ACE2 significantly influence the entry efficiency of the Omicron BA.5 variant (27). In our infectivity assay (**Figure 3E**), the N31D mutation in *R. blasii* ACE2 completely abrogated infectivity by BM48-31. Notably, the same N31 residue is also present in *Rhinolophus alcyone*, a species previously shown to be susceptible to BM48-31. These findings strongly support the role of N31 as a key determinant of S-ACE2 interaction. Furthermore, when both residues at positions 31 and 41 were simultaneously mutated in *R. blasii* ACE2, infectivity by RhGB01 was markedly reduced. In contrast, mutation of either position alone did not abolish viral entry. This pattern suggests a potential functional interdependence between the two residues. It is possible that residues 31 and 41 act cooperatively to stabilize S-ACE2 binding, and that compensatory structural effects from one residue may mask the loss of interaction from the other. Only simultaneous disruption appears sufficient to break the interaction network. These findings highlight the critical roles of residues 31 and 41—particularly position 31—as functionally conserved hotspots in determining sarbecovirus host range across different bat species.

To further investigate the molecular interactions underlying the S-ACE2 binding that contribute to differences in viral infectivity, we attempted to model the structure of the novel ACE2 variants identified in this study. However, due to limitations in current structure prediction tools, none of the available models reliably captured the structural features of our novel ACE2 sequences. As an alternative, we referred to a recently published cryo-electron microscopy structure describing the complex formed between the S protein of a clade 3 sarbecovirus, BtKY72, and the ACE2 receptor of an African bat, *Rhinolophus landeri* (23). In this reference structure, the S residues K482 and Y478 of BtKY72 are shown to interact directly with position 31 of ACE2. Notably, the BM48-31 S protein contains an alanine residue at the position corresponding to BtKY72 K482. This substitution from a positively charged lysine to a small hydrophobic alanine may shift the nature of the interaction at the ACE2 interface. In *R. blasii* ACE2, residue 31 is an asparagine (N31), which could form favorable interactions with both the hydrophobic alanine and the adjacent tyrosine. We hypothesize that this interaction network stabilizes S-ACE2 binding and facilitates viral entry. These structural insights, although extrapolated from a related species, support our functional findings that N31 in *R. blasii* ACE2 is critical for BM48-31 infectivity. They also underscore the importance of position 31 as a conserved contact point in clade 3 sarbecovirus-ACE2 complexes.

Overall, in this study, we report the ACE2 sequences of two African *Rhinolophus* bat species for the first time. Using a pseudovirus infection system, we demonstrated distinct infectivity profiles of various sarbecoviruses across cells expressing these novel ACE2 variants. Our findings highlight the critical role of ACE2 residues 31 and 41 in modulating S-ACE2 interactions. In particular, residue 31 appears to be a functional hotspot that may significantly influence sarbecovirus host range across species.

### Limitations of the study

While our findings provide new insights into ACE2 diversity and sarbecovirus infectivity, several limitations remain. First, only eight members of the sarbecovirus subgenus were included in the infectivity assays. Due to the geographical origins of the bat individuals used in this study, the majority of selected viruses belonged to clade 3, which are primarily found in Europe and Africa. As a result, the susceptibility of the novel ACE2 variants to clade 1 sarbecoviruses was not fully evaluated. Second, we employed a pseudovirus infection system rather than live virus assays. Although pseudovirus systems are widely used for studying viral entry mechanisms, they do not fully replicate natural infection conditions, as other viral proteins beyond the S protein may also influence infectivity. However, it should be noted that many clade 3 live viruses including BM48-31 and RhGB01 are not currently available to the best of our knowledge. Third, despite the novelty of the ACE2 sequences identified, the relatively small sample size limits the generalizability of our conclusions. In particular, only one *R. blasii* individual was analyzed. Although BM48-31 was originally detected in *R. blasii*, the viral sequence was isolated in Bulgaria, whereas our *R. blasii* sample originated from Zambia. Given this geographic separation, we cannot exclude the possibility that ACE2 in *R. blasii* is polymorphic, as observed in *R. simulator* in this study and in *Rhinolophus affinis* in previous studies (20, 21). Additional sampling of *R. blasii* from diverse regions will be necessary to fully understand ACE2 variability and its implications for virus susceptibility.

## Materials and Methods

### Ethics statement for bat sampling

Bat samples were collected as a part of the research project Molecular and Serological Surveillance of Viral Zoonoses in Zambia (DNPW8/27/1), approved by the Department of National Parks and Wildlife, Ministry of Tourism and Arts of the Republic of Zambia (act no. 14 of 2015).

### Sample collection and processing

Bats were collected using harp trap sampling. Animals were euthanized by overdose of Isoflurane. Samples of lung were taken from each body and were stored separately at – 80°C.

### Geographical mapping of sampling sites

African map and GIS border data were downloaded from SimpleMaps.com with a CC BY 4.0 license. GPS data of sampling sites (Table 2) were visualized on the map using QGIS (28)

**Table 2.** Sampling information.

| Sample ID | location | Sampling date | GPS |
| --- | --- | --- | --- |
| Z-Rsi-01 to Z-Rsi-04 | Shimabala, Zambia | 26.04.2016 | 15.647059° S,<br>28.232882°E |
| Z-Rbla | Suesueman, Zambia | 07.12.2016 | 15.601892° S,<br>28.724414° E |

### ACE2 sequencing from bat organs

For each specimen, the bat lung was homogenized by a homogenizer pestle. Total RNA was extracted using the QIAamp RNA Blood Mini Kit (Qiagen, Cat# 52304) according to the manufacturer’s instructions. Extracted RNA was reverse transcribed to cDNA using SuperScript III Reverse Transcriptase (Thermo Fisher, Cat# 18080085), following the supplier’s protocols using the specific primer designed for ACE2 (cDNA_bat-ACE2: 5’-ATTTACATACAATRAAATCACCTC-3’). The resulting cDNA served as the template for two rounds of PCR by same primer set (F-ACE2-out: 5’-AATGGGGTTTTGGCGCTCAG-3’, R-ACE2-out: 5’-CATACAATGAAATCACCTCAAGAG-3’) (29) performed with PrimeSTAR GXL DNA Polymerase (Takara Bio, Cat# R050A). Target bands (*Rhinolophus* ACE2 DNA) were excised, and DNA was purified with a gel extraction kit (Qiagen, Cat# 28704). Purified ACE2 DNA amplicons from each sample were subjected to Sanger sequencing (Eurofins) by following primers (bat-ACE2-Seq1: 5’-ATGTCAGGCTCTTCCTGGCT-3’, bat-ACE2-Seq2: 5’-GAGGTCGGCAAGCAGCTGAG-3’, bat-ACE2-Seq3: 5’-TTCAGGATCAAGATGTGCAC-3’, bat-ACE2-Seq4: 5’-GACCATTTTTGAATTCCAGTTTC-3’, bat-ACE2-Seq5: 5’-GAAGCCAAGAATCTCCTTCA-3’, bat-ACE2-Seq6: 5’-CACTCTGCTGAAGGATCTGC-3’). Resulting sequence fragments were assembled with SnapGene v8.1.1 (SnapGene software, (2–7)).

### RNA sequencing of lung samples

Total RNA from bat lung was used for RNA sequencing. cDNA library was subsequently synthesized using Illumina Stranded Total RNA Prep with Ribo-Zero Plus (Illumina, Cat# 20040525) following the manufacturer’s protocols. The library was then sequenced using the Illumina NovaSeq 6000 sequencing platform.

### Reconstruction of CYTB sequences from RNA-Seq data

First, RNA-Seq reads were trimmed using fastp (v0.21.0) with default parameters. The trimmed reads were then assembled into contigs using Trinity (v2.13.2) with default settings. To identify contigs corresponding to the CYTB gene, a BLASTN search (v2.15.0) was performed using the assembled contigs as the database and the CYTB gene sequence of a closely related species, *Rhinolophus macrotis* (GenBank accession KX261914.1), as the query. The top-hit contig with the highest bit score was extracted using blastdbcmd (v2.15.0).

### ACE2 and CYTB molecular phylogenetic analysis

Since bat species annotations registered in NCBI may include species identifications based solely on morphological information, and may additionally be affected by inconsistent nomenclature or potential miss annotations by submitters, we established a homology search-based pipeline that does not rely on NCBI annotation metadata. First, we performed BLASTn v2.16.0 search against NCBI nt database using all the sequences determined in this study and our previous study (20) as query with the following options (-max_target_seqs 1000 -evalue 0.000001), resulting a total of 1610 unique entries. Then, we performed an initial phylogenetic analysis with FastTree v2.2.0 (30) and extracted sequences that placed together with our samples and used as a dataset for the input of our phylogenetic analysis. The maximum likelihood (ML) trees of nucleotide sequences of CTYB and ACE2 were constructed by the following procedures: multiple sequence alignment (MSA) was constructed using MAFFT v7.526 (31)with the default option. Alignment sites with low coverage were excluded using gappyout method of trimAl v1.5.rev0 (32). The ML tree was reconstructed using IQ-TREE 3 v3.0.1 (33) under the TN+F+R5 nucleotide substitution model (**Figure 1B**) or TIM3+F+I+G4 nucleotide substitution model (**Figure 1C**) found by ModelFinder (34) with 1000 ultrabootstrap analyses (35). All CTYB and ACE2 sequences used in molecular phylogenetic analysis were summarized in **Table S1**.

### Sarbecovirus molecular phylogenetic analysis

The maximum likelihood tree for the RBD-encoding nucleotide sequences was reconstructed by the following procedures: an MSA was constructed woth MUSCLE using Aliview version 1.30 with the default option (36); The ML tree was reconstructed using IQ-TREE 3 v3.0.1 (33) under the TVM+F+I+G4 nucleotide substitution model found by ModelFinder (34) with 1000 ultrabootstrap analyses (35). All S RBD sequences used in molecular phylogenetic analysis were summarized in **Table S2**.

### Nucleotide and amino acid sequences data collection

The S sequences of sarbecoviruses used in this study were derived from our previous work (22), with the exception of PDF-2370 and PRD-0038. The PDF-2370 and PRD-0038 S sequences were NCBI GenBank (Genbank accession ID: MT726044.1, MT726045.1). The metadata of ACE2 sequences and information of the sarbecoviruses used in this study are summarized in Table S1 and S2.

### Plasmid construction

A part of the plasmids expressing human codon-optimized S protein of sarbecoviruses were prepared in our previous study (22). Plasmids expressing codon optimized PDF-2370 and PRD-0038 S proteins were synthesized by a gene synthesis service (Fasmac). DNA fragments expressing *Rhinolophus blasii* and *simulator* ACE2 were amplified by PCR (pW_ACE2_fw: 5’-CTAGCCTCGAGGTTTGGATCCGCCACCATGTCAGGCTCTTCC-3’; pW_ACE2_rv: 5’-TTAAACACTAGTACGCGTCTACTTGTCATCGTCATCCTTGTAATCGATGTCATGATCTTTATAATCACCGTCATGGTCTTTGTAGTCAAACGAAGTCTGA-3’). Plasmids expressing derivatives of ACE2 protein of *Rhinolophus blasii* were subjected to site-directed overlap extension PCR (pW_blasii_K31D_fw: 5’-CAAGATATTTTTGGACGACTTTAACACTG-3’; pW_blasii_K31D_Rv: 5’-CAGTGTTAAAGTCGTCCAAAAATATCTTG-3’; pW_blasii_H41Y_fw: 5’-AGCCGAAAACCTGTCTTACCAAAGTTCAC-3’; pW_blasii_H41Y_Rv: 5’-GTGAACTTTGGTAAGACAGGTTTTCGGCT-3’). The resulting PCR fragments of ACE2 were cloned into the BamHI/MluI site of pWPI-MCS-zeo vector (37) with 3xFLAG tag at the C-terminus, using In-Fusion-HD Cloning Kit (Takara, Cat# Z9650N). The plasmid expressing S protein of PDF-2370 and PRD-0038 were digested and cloned into the KpnI/NotI site of backbone of pCAGGS vector using T4 DNA ligase (New England BioLabs, Cat# M0202L). Nucleotide sequences were determined by DNA sequencing services (Eurofins), and the sequence data were analyzed by SnapGene v8.2.1.

### Cell culture

LentiX-293T cells (a human embryonic kidney cell line; Takara, Cat# 632180) were maintained in Dulbecco’s modified Eagle’s medium (high glucose) (Sigma-Aldrich, Cat#6429-500ML) containing 10% fetal bovine serum and 1% penicillin-streptomycin (Sigma-Aldrich, Cat# P4333-100ML). HEK293 cells (a human embryonic kidney cell line; ATCC, CRL-1573) were maintained in Dulbecco’s modified Eagle’s medium (low glucose) (Sigma-Aldrich, Cat#6046-500ML) containing 10% fetal bovine serum and 1% penicillin-streptomycin (Sigma-Aldrich, Cat# P4333-100ML). HOS cells (a human osteosarcoma cell line; ATCC CRL-1543) that stably express TMPRSS2 and *Rhinolophus* bat ACE2 were generated as described in the below (see “generation of HOS-TMPRSS2 cells stably expressing ACE2 proteins” section) and maintained in Dulbecco’s modified Eagle’s medium (high glucose) (Sigma-Aldrich, Cat# 6429-500ML), zeocin (50 μg/mL; InvivoGen, Cat# ant-zn-1) and G418 (400 μg/mL; Nacalai Tesque, Cat# G8168-10ML). Cell lines were tested for mycoplasma contamination by CycleavePCR Mycoplasma Detection Kit (Takara, Cat# CY232).

### Generation of HOS-TMPRSS2 cell lines stably expressing bat ACE2 proteins

HOS-TMPRSS2 cells stably expressing human ACE2 protein were prepared as described in our previous study (22, 25, 37). To prepare lentiviral vectors expressing *R. simulator* and *Rhinolophus blasii* ACE2, LentiX-293T cells (500,000 cells) were transfected with 0.2 µg of pCMV-VSV-G-RSV-Rev, 0.9 µg of psPAX2-IN/HiBiT and 0.9 µg of respective pWPI-ACE2-zeo plasmid using TransIT-293 (Takara, Cat# MIR2704) following the manufacturer’s protocol. 48 hours post transfection, the supernatant containing lentivector particle was harvested and filtered using 0.45 µm filter (Millipore, Cat# SLHVR33RB). HOS-TMPRSS2 cells (100,000 cells) were then transduced with prepared lentiviral vector, and selected using zeocin (50 μg/mL; Invivogen, Cat# ant-zn-1) and G418 (400 μg/mL; Nacalai Tesque, Cat# 09380-44) from 48 hours post transduction for 14 days.

### Western blot

Western blot was performed as described in previous study (25). HOS-TMPRSS2 cells stably expressing ACE2 proteins (see “Generation of HOS-TMPRSS2 cell lines stably expressing bat ACE2 proteins” section) were harvested, washed and lysed in lysis buffer consisting of RIPA buffer (50mM Tris-HCI buffer [pH 7.6], 150 mM NaCl, 1% Nonidet P-40, 0,5% sodium deoxycholate, 0.1% SDS) and protease inhibitor (Nacalai Tesque, Cat# 03969-21). The lysates supernatants were diluted with 2x sample buffer [100mM tris-HCl (pH6.8), 4% SDS, 12% β-mercaptoethanol, 20% glycerol, 0,05% bromophenol blue]. Samples were boiled for 10 minutes, then 10uL of the samples were subjected to Western blot. For detection, the following antibodies were used: mouse anti-alpha-tubulin (TUBA) monoclonal antibody (clone DM1A, Sigma-Aldrich, Cat# T9026, 1:10000), Horseradish peroxidase (HRP)-conjugated mouse anti-FLAG monoclonal antibody (clone M2, Sigma-Aldrich, Cat# A8592, 1:1000), and HRP-conjugated horse anti-mouse IgG antibody (Cell Signaling, Cat # 7076S, 1:2000). Chemiluminescence was detected using Western Lightning Plus-ECL (PerkinElmer, Cat# NEL104001EA) according to the manufacturer’s instruction. Bands were visualized using ChemiDoc Touch Imaging System (Bio-Rad).

### Pseudovirus infectivity assay

Pseudovirus assay was performed as previously described (38–40). In brief, HIV-1 based, luciferase expressing reporter virues were pseudotyped with S proteins of sarbecoviruses. LentiX-293T cells (500,000 cells) were transfected with 0.8 μg psPAX2-IN/HiBiT (38), 0.8 μg pWPI-Luc2 (41), and 0.4 μg of plasmid expressing respective sarbecovirus S using TransIT-293 (Takara, Cat# MIR2704) following the manufacturer’s protocol.48 hours post-transfection, the culture supernatants were harvested and filtered using 0.45 µm filter (Millipore, Cat# SLHVR33RB). Filtered pseudoviruses were stored at −80 °C until use. Prior to pseudovirus infection, the amount of input virus was normalized to the HiBiT value measured by NanoGlo HiBiT lytic detection system (Promega, Cat# N3040) (41). The system relies on a HiBiT tag fused to the C-terminus of integrase within lentiviral particles. Upon interaction with LgBiT, functional NanoLuc luciferase is reconstituted, producing luminescence in the presence of substrate. In each pseudovirus particle, the detected HiBiT value is correlated with the amount of the pseudovirus capsid protein, HIV-1 p24 protein. Therefore, we calculated the amount of HIV-1 p24 capsid protein based on the HiBiT value measured, according to the previous paper (41). The amountof HIV-1 p24 antigen used in the assay was 4ng. HOS-TMPRSS2 cells stably expressing *R. simulator* and *blasii* ACE2 and HEK293 cells transfected with plasmids expressing *Rhinolophus blasii* and its derivatives with PEI-Max (Polysciences, Cat# 24765-1) were used for the assay. Cells were infected with pseudoviruses respectively, and infected cells were lysed with a Bright-Glo Luciferase-Assay System (Promega, Cat# E2620) 2 days post infection. The luminescent signal was measured using a GloMax Explorer Multimode Microplate Reader (Promega).

## DATA AVAILABILITY

The computational codes used in the present study are available on the GitHub repository (https://github.com/TheSatoLab/XXXXX).

All new ACE2 and CYTB sequences, and RNA-seq data described in this study will be deposited in GenBank database upon manuscript acceptance.

Any additional information required to reanalyze the data reported in this work is available from the lead contact upon request.

## Author Contributions

MK, AT, KC, BMH conducted sample collection;

SF performed RNA extraction and ACE2 sequencing;

YZ performed plasmid construction and virological experiments;

AK and YS performed RNA sequencing;

JI performed reconstruction of CYTB sequences from RNA-Seq data;

SL and SK constructed the Rhinolophus CYTB sequence dataset;

SF and YZ performed phylogenetic and phylogeographic analyses;

YZ, SF and KS designed the experiments and interpreted the results;

YZ, SF and KS wrote the original manuscript;

All authors reviewed and proofread the manuscript.

## Acknowledgments

We thank Dr. Kenzo Tokunaga (National Institute for Infectious Diseases, Japan) for providing reagents. We thank Dr. Luca Nishimura (The University of Tokyo, Japan) for support in the phylogenetic analysis. The super-computing resource was provided by Human Genome Center at The University of Tokyo.

This study was supported in part by AMED ASPIRE Program (25jf0126002, to Kei Sato); AMED SCARDA Japan Initiative for World-leading Vaccine Research and Development Centers “UTOPIA” (JP223fa627001, to Kei Sato, Jumpei Ito, Shusuke Kawakubo, Spyros Lytras); AMED SCARDA Program on R&D of new generation vaccine including new modality application (253fa727002, to Kei Sato); AMED Research Program on Emerging and Re-emerging Infectious Diseases (24fk0108907, JP25fk0108690, to Kei Sato); AMED Japan Program for Infectious Diseases Research and Infrastructure (Collaborative Research via Overseas Research Centers) (25wm0225041, to Kei Sato); JSPS Bilateral Program JPJSBP123456789 and JSPS-VAST Joint Research Program (120259601, to Kei Sato); JSPS KAKENHI Fund for the Promotion of Joint International Research (International Leading Research) (JP23K20041, to Kei Sato); JST PRESTO (JPMJPR22R1, to Jumpei Ito); JSPS KAKENHI Grant-in-Aid for Scientific Research B (JP25K00116, to Jumpei Ito); JSPS Research Fellow DC2 (24KJ0628, to Shigeru Fujita); JSPS KAKENHI Grant-in-Aid for Scientific Research A (JP24H00607, to Kei Sato); The Cooperative Research Program (Joint Usage/Research Center program) of Institute for Life and Medical Sciences, Kyoto University (to Kei Sato); Mitsubishi UFJ Financial Group, Inc. Vaccine Development Grant (to Kei Sato); SHIONOGI Infectious Disease Research Promotion Foundation (to Jumpei Ito); AIST KAKUSEI project (FY2024 and FY2025 to Shusuke Kawakubo).

